# Prolonged Single Neuron Voltage Imaging in Behaving Mammals

**DOI:** 10.1101/2025.05.29.656886

**Authors:** Yangyang Wang, Cara Ravasio, Yuxin Zhou, Xue Han

**Author notes:** These authors contributed equally to this work (Yangyang Wang, Cara Ravasio).

## Abstract

Long-term recording of single-neuron membrane voltage dynamics during behavior is highly desirable. Using ElectraOFF, a fully genetically encoded, photostable, high-performance fluorescence voltage indicator, we achieve routine cellular-resolution imaging over tens of minutes, occasionally up to eighty minutes, in behaving mice with minimal signal loss across neuron types and brain regions. This extended recording capability reveals plasticity changes throughout intracranial electrical neuromodulation, highlighting the novel insights enabled by prolonged voltage imaging.

## Main Text

The new generation of genetically encoded voltage indicators (GEVIs) enables high spatiotemporal resolution imaging of membrane voltage (Vm) from individual neurons in the mammalian brain^1–9^. In addition to capturing spiking, GEVIs can detect subthreshold dynamics from genetically defined neurons, which has not been possible with traditional extracellular recording or calcium imaging methods^10–13^. However, most GEVIs still suffer from poor photostability, generally limiting recording duration to about 10-20 minutes^1–9^, but success for longer recordings lasting tens of minutes has been demonstrated with GEVI JEDI-2P (∼40 mins)^7^ and ASAP5 (∼1 hour)^6^ using two-photon microscopy. The newly developed green GEVI ElectraOFF^14^, exhibits high voltage sensitivity, excellent photostability, and is compatible with a widefield microscope that permits simultaneous high-speed recording of many cells. Furthermore, ElectraOFF is fully genetically encoded, without needing any chemical cofactors, greatly improving its applicability *in vivo*. We tested the performance of ElectraOFF in different types of neurons across the hippocampal CA1, visual cortex and striatum in voluntarily locomoting mice, and used ElectraOFF to probe cellular plasticity during chronic electrical stimulation widely used in clinical neuromodulation.

To characterize the performance of ElectraOFF in the brain of awake mice, we transduced hippocampal CA1 pyramidal neurons with AAV9-CaMKII-ElectraOFF and installed a glass cranial window for chronic optical access (see **Methods**). ElectraOFF-expressing neurons were imaged using a custom widefield fluorescence microscope at a frame rate of 667 Hz, while mice were head-fixed and voluntarily locomoting. Image frames were collected for 30 s every minute to allow for data streaming, and the recorded image stacks were motion corrected and processed offline to obtain the Vm traces and spike timing for each neuron. We started imaging with a low LED power at ∼20 mW/mm^2^ to minimize photobleaching and gradually increased the power every 5-20 min to compensate for the loss of fluorescence (up to ∼200 mW/mm^2^). Using this experimental protocol, we were able to obtain recordings that lasted for up to 80 min without compromising spike quality (**Fig. 1a-c and Fig. S1**).

**Figure 1.**
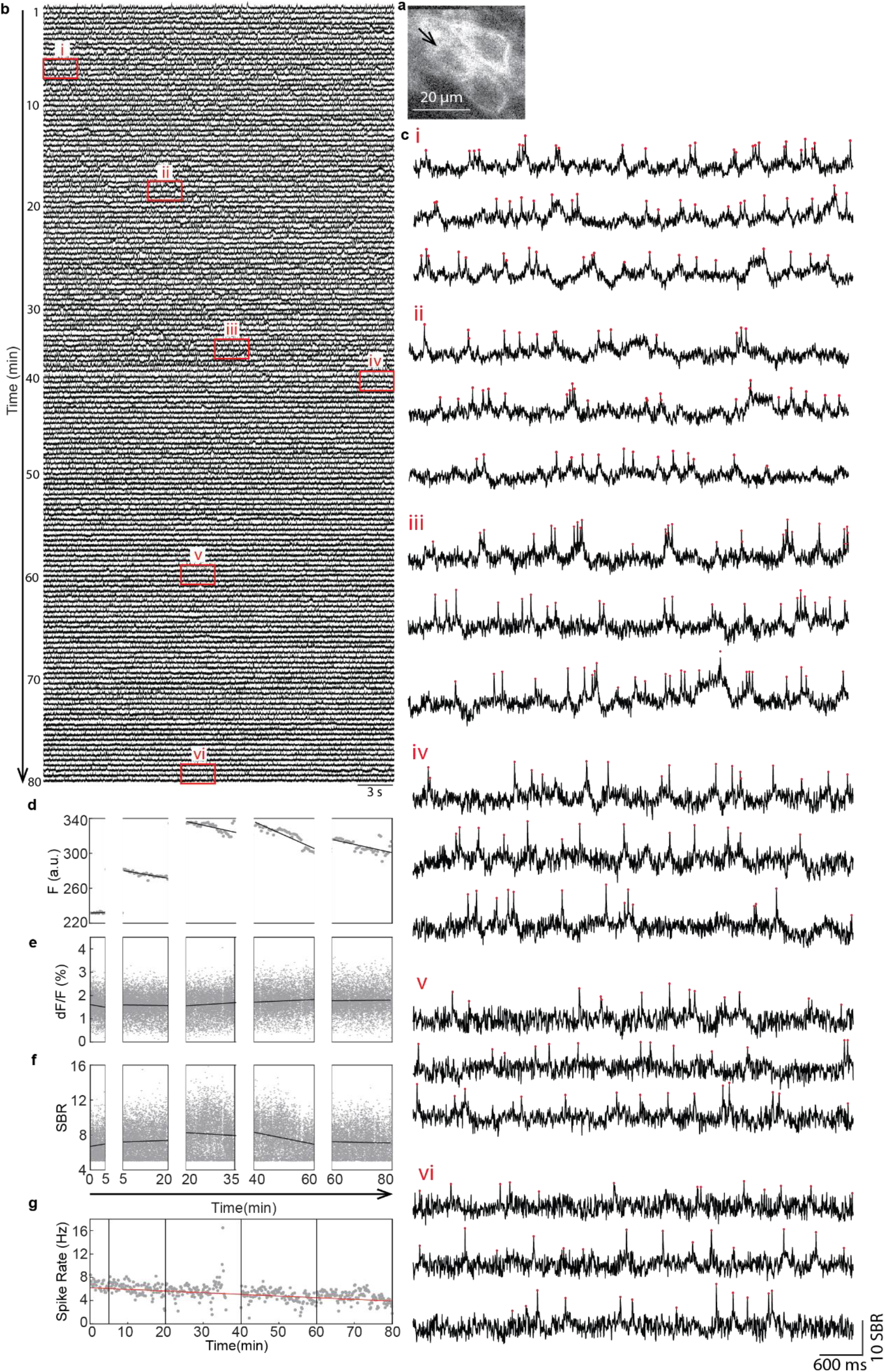
An example 80-minute-long voltage recording from a CA1 neuron in an awake mouse. **a**) An example imaging field of view showing three CA1 neurons expressing ElectraOFF. Shown is the maximum-minus-minimum fluorescence projection image, with the black arrow indicating the example neuron whose Vm (black traces) and spikes (red dots) are displayed. **b**) Vm recorded throughout the 80-minute-long duration. **c**) Zoomed-in view of periods highlighted by the corresponding boxes in b). **d**) Electra raw fluorescence intensity (F) over time. LED power was incrementally increased to compensate for photobleaching and is indicated by gaps in the x-axis. Each dot represents the averaged raw F within each 30 s trial. The black lines indicate linear fit within each time period with constant LED power. **e**) Similar to d), but for normalized spike amplitude (dF/F). Each dot corresponds to a spike, and the linear fit is shown in black. **f**) Similar to e), but for spike SBR. **g**) Spike rate over time. LED power shifts are indicated by vertical black lines and the red line is the linear fit across the entire recording. Each dot corresponds to a 10 s bin of the recording.

To quantify ElectraOFF’s photostability, we computed raw Vm fluorescence (F), normalized spike amplitude (dF/F; with dF defined as the fluorescence difference between the spike peak and the minimum value within the preceding 5 ms), spike signal-to-baseline ratio (SBR; defined as spike dF divided by variance of spike-removed Vm fluctuations during each 30-second trial), and spike rate (defined as the mean rate in each 10-second bin; see **Methods**). Photobleaching rates were estimated by linearly fitting across all data collected at the same LED power, except spike rate which was fit across the entire recording because little relationship was detected between LED power and spike rate (**Fig. S2m**). We found that for the example neuron shown in **Fig. 1a-c**, raw F declined at 0.21 ± 0.18% per minute (mean ± s.d. here and throughout unless otherwise stated; n=5 testing periods with varying LED power) throughout the 80 min recording time (**Fig. 1d**), spike rate decreased at 0.5% per minute (R^2^=0.14, p <0.001) (**Fig. 1g**), while spike dF/F, and SBR remained remarkably stable with decay rates of 0.12 ± 0.76 and 0.04 ± 0.61% per minute respectively (**Fig. 1e, f and Fig. S2a**). Thus, while ElectraOFF exhibits some photobleaching of raw F and a decrease in detectable spikes, compensatory LED adjustments can maintain stable spike dF/F and SBR, ensuring spike quality and enabling reliable long-term Vm recordings in CA1 of awake animals.

To systematically evaluate the performance of ElectraOFF across cell types and brain regions in behaving mice, we expressed ElectraOFF in striatal cholinergic (ChI) neurons of Chat-Cre mice and in visual cortical parvalbumin (PV) interneurons of PV-Cre mice using AAV9-CAG-DIO-ElectraOFF. ElectraOFF supported stable voltage imaging that routinely lasted for about 10 min in striatal ChIs and for tens of minutes in cortical PVs (**Fig 2a, b, n**). For the three example neurons, the bursting ChI, the tonically firing ChI, and the PV neuron, we detected minimal photobleaching of raw F at 2.9 ± 1.0%, 2.3 ± 5.0% and 1.3 ± 0.55% per minute respectively (**Fig. 2c, o, Fig. S2e**), moderate decay in spike rate at 7.2%, 8.9%, and 2.6% per minute (R^2^=0.78, p<0.001; R^2^=0.52, p<0.001; R^2^=0.23, p<0.001), and negligible decay in spike dF/F, and spike SBR (**Fig. 2d-f, p-r and Fig. S2f-h**). Across the population of recorded CA1 pyramidal neurons, striatal ChIs and cortical PVs, raw F photobleached at 0.72 ± 0.30%, 1.3 ± 0.89%, 2.7 ± 2.7% per minute respectively (**Fig. 2t and Fig. S2i**), and spike rate decayed at 3.4 ± 2.4%, 8.5 ± 1.7%, 5.0 ± 6.2% per minute respectively (**Fig. 2w, Fig.S2j**), while dF/ F, spike SBR showed little decay (**Fig. 2u, v and Fig. S2k,l**). Furthermore, ElectraOFF reliably distinguished striatal ChIs exhibiting spike bursts (**Fig. 2g**) versus tonic firing (**Fig. 2h)**. Aligning Vm to spikes revealed prominent Vm delta oscillation in bursting ChIs (**Fig. 2i, k**), but not in tonically firing ChIs (**Fig. 2j, l**), underscoring ElectraOFF’s ability to detect subthreshold Vm dynamics beyond spike timing.

**Figure 2.**
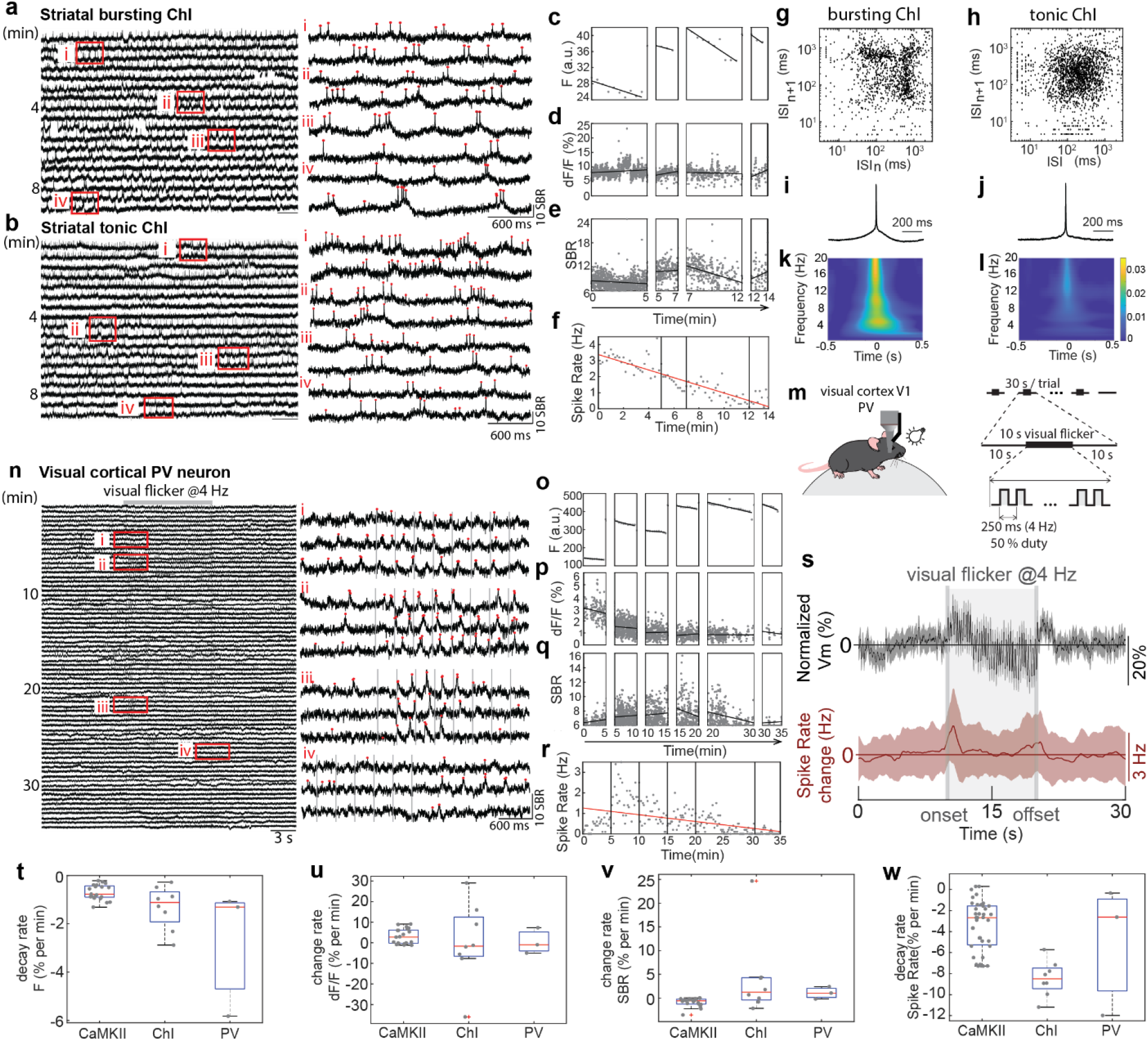
ElectraOFF is photostable across cell types and brain regions during both intrinsic and evoked activity. **a**) left, Vm from an example striatal cholinergic interneuron (ChI) with bursty firing in a voluntarily locomoting mouse. right, zoomed-in views of Vm for the periods i-iv shown on the left. **b**) Similar to a, but for a ChI with tonic firing. **c**) Raw F of the example bursting ChI over 14 min of the recording. The adjustments of LED power are indicated by gaps. Each dot represents the mean raw F of a 30 s trial. The black lines are linear fits with periods having constant LED power. **d**) Similar to c), but for normalized spike dF/F. Each dot corresponds to a spike. **e**) Similar to d), but for spike SBR. **f)** Spike rate over time. LED power shifts are indicated by vertical black lines and the red line is the linear fit across the entire recording. Each dot corresponds to a 10 s bin of the recording. **g**) The inter-spike interval (ISI) return map of all spikes recorded for the example bursting ChI shown in a) (n = 1411 spikes). **h**) Similar to g), but for the example tonic ChI shown in b) (n = 1203 spikes). **i**) Spike-triggered Vm for example bursting ChI. Black line is the mean, and shaded areas indicate 95% confidence interval. **j**) Similar to i), but for the example tonic ChI. **k**) Spike-triggered Vm power spectrogram for the bursting ChI. **l**) Similar to k), but for the tonic ChI. **m**) Experimental schematic for visual flicker presentation to a head-fixed voluntarily locomoting mouse, while recording visual cortical PV interneurons. Visual flickers were delivered at 4 Hz, for 10 s during each 30 s trial. **n**) Vm traces of an example PV neuron across 35 min recording. **o-r**) Similar to c-f), but for the example PV. **s**) Vm (black) and spike rate (red) changes aligned to visual flicker onset. Shading areas indicate standard error of mean (n = 70 trials). **t**) The photobleaching rates of raw F for CA1 neurons (n=21 neurons), striatal ChIs (n=8 neurons), and visual cortical PV neurons (n=3 neurons). Each dot represents a neuron. Red lines are the mean, and the boxes mark the interquartile range. **u-w**) Similar to t), but for spike dF/F, SBR, and spike rate respectively (n=21 for CA1 neurons, n=8 for ChIs, and n=3 for PVs).

To test the ability of ElectraOFF in capturing behaviorally relevant stimuli, we presented mice with 4 Hz visual flickers for 10 s in the middle of each 30-second-long imaging trial (**Fig. 2m, n**). We found that visual flicker evoked a transient Vm depolarization in visual cortical PV neurons that lasted for about 1 s, which was followed by sustained hyperpolarization throughout the stimulation period and a pronounced rebound after stimulation offset (**Fig. 2s**). Interestingly, while spiking increased during the initial Vm depolarization, spike rate quickly returned to pre-stimulation levels even though Vm remained hyperpolarized, highlighting ElectraOFF’s ability to capture long-lasting synaptic inhibition during behavior.

Finally, we investigated the sustained effects of intracranial electrical stimulation on CA1 neurons to understand the impact of prolonged stimulation on cellular plasticity. We delivered biphasic square-wave pulses (100 µs/phase) at 40 Hz that have been shown to entrain CA1 neurons^15^. Stimulation was delivered locally for 1 min in the middle of each 90 s trial (**Fig. 3a**) at a current intensity sufficient to evoke a neural response without inducing detectable behavioral changes (**Methods**). Each neuron was tested for 5 trials with an intertrial interval of 1.5 min. Consistent with previous observations, of the 49 neurons recorded, 27 (55%) were significantly entrained (**Fig. 3f**), exhibiting greater Vm spectral power at 40 Hz during stimulation than baseline. Across all recorded neurons, stimulation did not change population firing rate, but it evoked a pronounced Vm hyperpolarization immediately after stimulation onset that lasted for ∼1 s, which was followed by Vm depolarization that stabilized after ∼15 s (**Fig. 3b**). To further evaluate the time-course of the evoked responses, we divided the stimulation period into four consecutive 15-second epochs and compared the 40 Hz spectral power of Vm across epochs for every trial (**Fig. 3c**). We found that entrainment strength across the entrained neurons declined overtime within each trial, with the highest 40 Hz power observed during the first 15 s (**Fig. 3d, e**). Furthermore, we detected a gradual reduction in entrainment strength across successive trials (**Fig. 3g**). Thus, prolonged electrical stimulation-evoked Vm changes progressively decay over time, underscoring the capacity of ElectraOFF to uncover plasticity mechanisms in neural circuits.

**Figure 3.**
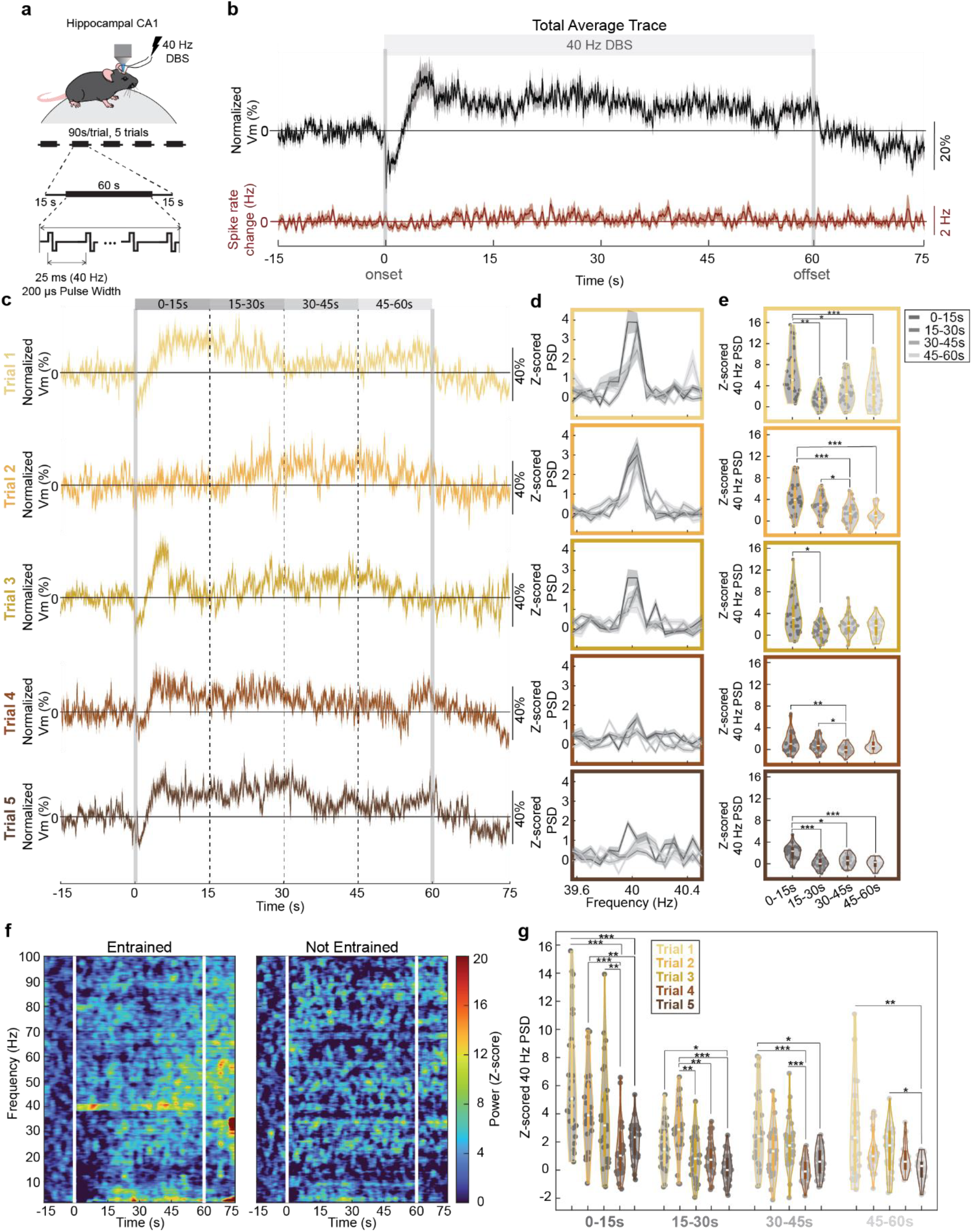
Sustained ElectraOFF recordings revealed intracranial electrical stimulation induced cellular plasticity effects. **a**) Top, schematic of the experimental setup. Bottom, stimulation protocols. Stimulation was delivered locally in CA1, while recording Vm of CA1 pyramidal neurons. Each field of view underwent five stimulation trials with a 1.5 min inter-trial interval. Each trial included 15 s pre-stimulation baseline, 60 s electrical stimulation (40 Hz, biphasic pulses), and 15 s post-stimulation periods. **b**) Stimulation evoked change in Vm (black) and spike rate change (red). Vm change is normalized to mean spike height and shifted by the mean pre-stimulation baseline Vm. Shaded areas represent standard error of mean (SEM) (n = 49 neurons). **c**) Trial-wise average Vm traces with SEM shown as shading (n=49). The stimulation periods were subdivided into 15-second epochs as highlighted on top). Vertical lines denote stimulation onset and offset (solid) and 15-second subdivisions (dotted). **d**) Power spectral density (PSD) of Vm traces, computed for each 15-second epoch and normalized to the baseline PSD for each trial. Gray lines indicate mean PSD across neurons; shaded regions indicate SEM (n=49). **e**) Violin plots of 40 Hz PSD across stimulation epochs for entrained neurons (n=27) (Statistical details in Supplemental Table 1). **f**) Vm spectrograms of entrained (left, n=27) and not-entrained (right, n=22) neurons aligned to stimulation onset. **g**) Similar to e) but compared across trials for each epoch. (Statistical details in Supplemental Table 1). Violin plots depict the kernel density overlaid with box plots showing the interquartile range (1x, 1.5x). The white lines in the boxes are the median. All tests are two-tailed. *p < 0.05, **p < 0.01, ***p < 0.005.

## Discussion

ElectraOFF exhibits minimal photobleaching in the brains of behaving mice, with raw fluorescence intensity decaying at ∼0.5-1.3% per minute. This modest reduction allows for routine imaging for tens of minutes across cell types and brain regions. By progressively increasing LED power over time during prolonged imaging to minimize excitation light exposure, we were able to image most neurons for tens of minutes, and up to 80 minutes in CA1, without compromising spike quality. As spike detection depends on spike peak fluorescence, we noted a spike rate decay at a low rate of 6% per minute, while maintaining spike dF/F and spike SBR. Photobleaching is intrinsic to fluorescence imaging, and it is important to include control analysis when examining long term changes to rule out photobleaching effects. Voltage-dependent ElectraOFF fluorescence can be readily recorded with a high-speed widefield microscope, allowing for simultaneous recording from multiple neurons which cannot be easily achieved with high-speed two-photon or confocal microscopy techniques that have limited field-of-view sizes^9,16,17^.

Using ElectraOFF, we captured prominent Vm hyperpolarization upon visual flicker presentation in visual cortical interneurons and upon electrical stimulation in CA1 pyramidal neurons. These results highlight the unique advantage of voltage imaging in detecting synaptic inhibition, not possible using conventional extracellular electrodes or calcium imaging. As Vm determines spike timing, the ability to record Vm over an extended time unlocks new frontiers for analyzing the subthreshold decision-making processes of individual neurons during behavior and pathology. As an example, we demonstrated that upon sustained stimulation, either through natural visual flicker presentation or artificial intracranial electrical current delivery, Vm exhibits complex temporal changes, switching between depolarization and hyperpolarization which was often not accompanied by spiking changes. In particular, during chronic intracranial electrical stimulation, evoked Vm entrainment progressively decayed over time, revealing robust plasticity mechanisms related to clinical electrical neuromodulation.

**Supplementary Table 1:** Statistical Details for Fig. 3.

| **Stats Table** |  |  |  |  |  |
| --- | --- | --- | --- | --- | --- |
| Fig 3e p-values (Friedman test with Bonferroni multiple comparisons correction) |  |  |  |  |  |
|  | Trial 1 | Trial 2 | Trial 3 | Trial 4 | Trial 5 |
| 0-15s vs 15-30s | $4.36 \times 10^{-6}$ | 0.161 | 0.027 | 1 | $1.26 \times 10^{-5}$ |
| 0-15s vs 30-45s | 0.009 | $4.36 \times 10^{-6}$ | 0.122 | 0.001 | 0.013 |
| 0-15s vs 45-60s | $9.27 \times 10^{-5}$ | $8.16 \times 10^{-7}$ | 0.161 | 0.271 | $2.52 \times 10^{-6}$ |
| 15-30s vs 30-45s | 0.439 | 0.037 | 1 | 0.027 | 0.550 |
| 15-30s vs 45-60s | 1 | 0.013 | 1 | 1 | 1 |
| 30-45s vs 45-60s | 1 | 1 | 1 | 0.550 | 0.271 |
| Fig 3g p-values (Friedman test with Bonferroni multiple comparisons correction) |  |  |  |  |  |
|  | 0-15s | 15-30s | 30-45s | 45-60s |  |
| Trial 1 vs Trial 2 | 1 | 0.160 | 0.707 | 1 |  |
| Trial 1 vs Trial 3 | 0.478 | 1 | 1 | 1 |  |
| Trial 1 vs Trial 4 | $8.10 \times 10^{-8}$ | 1 | $5.08 \times 10^{-5}$ | 0.707 | |
| Trial 1 vs Trial 5 | $5.08 \times 10^{-5}$ | 0.020 | 0.008 | 0.002 | |
| Trial 2 vs Trial 3 | 1 | 0.002 | 1 | 1 |  |
| Trial 2 vs Trial 4 | $2.21 \times 10^{-5}$ | 0.001 | 0.059 | 1 | |
| Trial 2 vs Trial 5 | 0.004 | $3.62 \times 10^{-7}$ | 1 | 0.059 | |
| Trial 3 vs Trial 4 | 0.002 | 1 | $7.52 \times 10^{-4}$ | 1 | |
| Trial 3 vs Trial 5 | 0.098 | 0.707 | 0.059 | 0.020 |  |
| Trial 4 vs Trial 5 | 1 | 1 | 1 | 0.478 |  |

**Figure S1.**
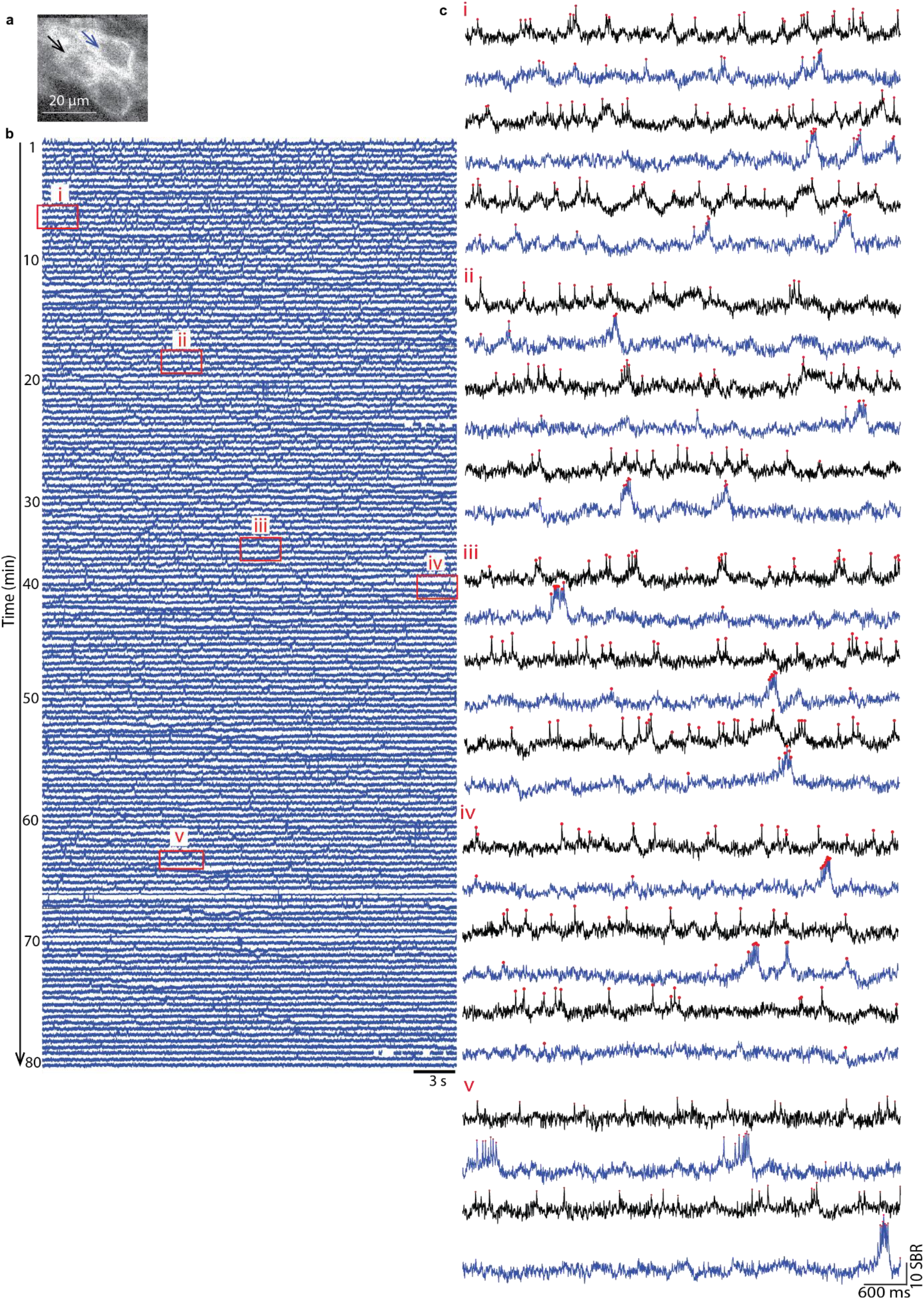
An example 80-minute long voltage recording from two adjacent CA1 neurons in an awake mouse. **a**) The same imaging field of view as in Fig. 1a, showing three neurons expressing ElectraOFF. The black arrow indicates the neuron whose Vm is shown in Fig. 1a, and the blue arrow indicates a neuron whose Vm is shown in b). Spikes are indicated by red dots. **b**) Vm throughout the 80-minute-long recording for the neuron indicated by the blue arrow in a. **c**) Zoomed in view of Vm from the two simultaneously recorded neurons during periods highlighted by the corresponding boxes in b). The blue and black traces are from the neurons indicated by the blue and black arrows in a) respectively.

**Figure S2.**
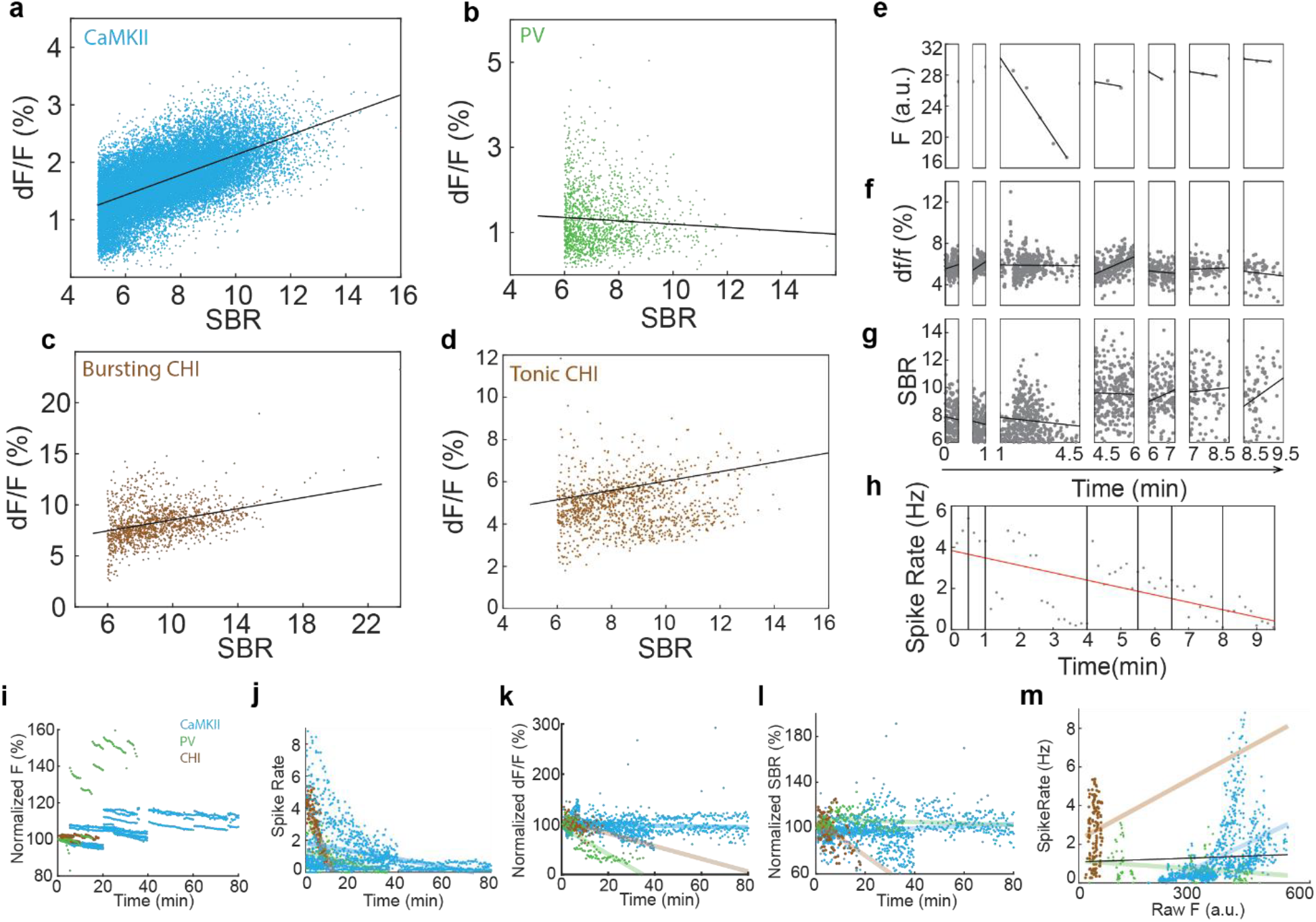
Detailed characterization of ElectraOFF photostability across cell types. **a**) The relationship between dF/F and spike SBR of the example CA1 neuron shown in Fig. 1a. Each dot corresponds to a spike. The black line indicates the linear fit (n=24365 spikes, R^2^=0.362, k=0.002, p < 0.001). **b-d**) Similar to a), but for the example PV neuron (n=1435 spikes, R^2^=0.004, k=-0 0004, p < 0.05), bursting ChI (n=1411 spikes, R^2^=0.122, k=0.003, p < 0.001), and tonic ChI (n=1208 spikes, R^2^=0.155, k = 0002, p<0.001) from Fig. 2. **e**) Raw F of the example tonic ChI from Fig. 2 over 14 min of the recording. The adjustments of LED power are indicated by gaps. Each dot represents the mean raw F of a 30 s trial. The black lines are linear fits with periods having constant LED power. **f)** Similar to e), but for normalized spike dF/F. Each dot corresponds to a spike. **g)** Similar to f), but for spike SBR. **h)** Spike rate over time. LED power shifts are indicated by vertical black lines and the red line is the linear fit across the entire recording. Each dot corresponds to a 10 s bin of the recording. **i)** The average raw F over time for all neurons. Each dot corresponds to the mean for each 30 s long recording period normalized to the first point of the given neuron. Cell type is color coded (Blue: 21 CA1 neurons (n=1174 recording periods); Green: 3 visual cortical PV neurons (n=120 recording periods); Brown: 8 striatal ChIs (n=126 recording periods). **j-l**) Similar to i), but for the spike rate, normalized dF/F, and SBR. The colored lines indicate the linear fit trendline for each cell type. **m)** Similar to j) but spike rate vs raw F (proxy for LED power). The black line is the linear fit across all neurons regardless of cell type and indicates there is a very weak correlation between the spike rate and raw F (n = 1420, k=0.007, R^2^=0.003, p=0.03).

## Methods

### Mouse Surgical Preparation

All animal experiments were performed in accordance with the National Institute of Health Guide for Laboratory Animals and approved by the Boston University Institutional Animal Care and Use and Biosafety Committees. Same-sex littermates were housed together prior to surgery and singly housed post-surgery. Cages were enriched with igloos or running wheels. Animal facilities were maintained around 70° F and 50% humidity and mice were kept on a 12 h light/dark cycle. A total of 3 adult mice, 1 ChAT-Cre female, 1 PV-Cre male (JAX stock #017320, B6.129P2-Pvalb^tm1(cre)Arbr^/J, Jackson Laboratory), and 1 C57BL/6 male (Charles River Laboratories Inc.), 16-24 weeks old at the start of the experiment were used in this study.

Under isoflurane general anesthesia, mice were surgically implanted with a sterilized recording ydevice for hippocampal or striatal recordings, or a glass coverslip for cortical recordings. Mice were provided with sustained release (SR) buprenorphine (3.25 mg/kg) pre-operatively that provides 72 hours of analgesia. Mice were singly housed to avoid damage to the recording apparatus after surgery.

For CA1 recordings, the recording device consisted of a custom imaging window attached to a guide infusion cannula and two stimulation electrodes. For striatal recordings, the recording device consisted of a custom imaging window attached to a guide infusion cannula. The imaging window consisted of a stainless-steel cannula (outer diameter 0.317 cm, inner diameter 0.236 cm, height 1.75 mm for CA1 window and 2 mm for striatal window; Item 3ADD3; Grainger), adhered to a circular coverslip (size 0, outer diameter 3 mm; No. CS-3R-0; Warner Instruments) using an ultraviolet-curable adhesive (Norland Products). The infusion guide cannula (8 mm, 26 gauge; P1 Technologies) was soldered at a 60° angle relative to the coverslip with the end flush to the base of the imaging cannula. The two stimulation electrodes (polyamide-coated steel core wire, diameter 127 µm, No. 7N003736501F; P1) were soldered onto dip pins (No. ED85100-ND; DigiKey), and secured to the infusion cannula with adhesive gel (Loctite 454; Loctite SF 713). The electrodes were bent so that the male end of the dip pin was perpendicular to the infusion cannula, while the electrode wire was parallel to the infusion cannula. The tips of the electrodes were about 3 mm apart and terminated below the imaging window. The recording apparatus was implanted over CA1 (centered at the stereotaxic coordinates: AP −2.0 mm, ML 1.7 mm, DV −1.5 mm from pia), or over the striatum (centered at the stereotaxic coordinates: AP 0.5 mm, ML 1.8 mm, DV −2 mm from pia) after gently aspirating the overlying cortical tissue and thinning the corpus callosum. A custom aluminum head-plate was attached to the skull posterior to the window via dental cement (Parkell, S380 and Stoelting, 5145). After a full recovery from the surgery (about 10 days), mice with a recording apparatus were infused with 1.5 µL of AAV9-CaMKII-ElectraOFF or AAV9-CAG-DIO-ElectraOFF (>10^13^ GC/mL, Shanghai Sunbio Medical Biotechnology and Charles River Laboratories), at a speed of 200 nL per minute, using a 10 µL syringe (World Precision Instruments, NANOFIL 10 µL syringe) controlled by a micro-infusion pump (Micro-2T, World Precision Instruments) attached to 28 gauge PTFE tubing (No. STT-28; Component Supply Co.) with a 33 gauge infusion cannula (8 mm with 1 mm projection; P1 Technologies). The infusion cannula was removed 10 min after the infusion had been completed to improve AAV spread.

For cortical recordings, a small craniotomy was created over the visual cortex (centered at the stereotactic coordinate: AP −3.6 mm, ML 2.3 mm), and a total of 1.5 µL of AAV9-CAG-DIO-ElectraOFF was injected via a 10 µL syringe (World Precision Instruments, NANOFIL 10 µL syringe) controlled by a micro-infusion pump (MicroPump4, World Precision Instruments) in 3 locations within the craniotomy, avoiding major blood vessels, ∼150 µm below pia. After viral injection the syringe was left in place for 10 min at each location, and then the syringe was removed. A circular glass coverslip (size 0, outer diameter 3 mm; No. CS-3R-0; Warner Instruments) was then pressed onto the cortex to ensure stable optical recordings and was secured to the skull with ultraviolet-curable cement (Tetric EvoFlow; Ivoclar). A custom aluminum head-plate was attached to the skull posterior to the window via dental cement (Parkell, S380 and Stoelting, 5145).

### Voltage Imaging

Animals were imaged starting 2 weeks post viral infusion in the CA1, 3 weeks in the striatum, and 4 weeks in the visual cortex, to ensure sufficient ElectraOFF expression. ElectraOFF voltage imaging was performed with a custom wide-field fluorescence microscope equipped with a sCMOS camera (Hamamatsu Photonics, C13440-20CU) and a 40X objective (Nikon, CFI Apo NIR 40×/0.8 NA W). A blue 470 nm LED (Thorlabs, M470L4) controlled by a T-Cube LED driver (LEDD1B, low gain; Thorlabs Inc.) was sequentially coupled to a 505 nm long-pass filter (Thorlabs, DMLP505R), excitation filter (FF01-390/482/532/640-25), quadband dichromatic mirror (Semrock Di03-R405/488/532/635), a multi-band dichromatic mirror (Chroma Technology Corp., ZT405/514/635rpc), and a 525 ± 22.5 nm bandpass emission filter (Semrock, FF01-525/45) emission filter.

ElectraOFF fluorescence images were collected using HCImage Live (Hamamatsu Photonics), triggered via a TTL pulse generated by a NI DAQ (National Instruments, BNC-2110 and PCIe-6323) controlled by MATLAB. Similarly, the electrical stimulator or visual flicker LED, if applicable, was also triggered by TTL pulses. HCImage Live collected up to 90 s of data (20000 or 60000 frames) with an exposure time set at 1.5 ms. HCImage was set to capture 128×512 pixels with 2×2 binning, corresponding to 41×164 µm^2^. Each trial was saved in a separate DCAM file. The files were analyzed offline with MATLAB. Camera frame exposure from the Hamamatsu camera, electrical stimulation pulses, and visual flicker pulses were simultaneously recorded as TTL pulses with an Open Ephys acquisition board (Open Ephys, SKU: OEPS-9030) at a rate of 30 kHz to ensure proper alignment during further analysis offline^18^.

### Electrical Stimulation

Electrical stimulation was delivered by a constant current stimulator (Model 4100; A-M Systems). The stimulation waveform was a cathode-leading biphasic square wave pulse delivered at 40 Hz with a 200 µs total pulse width (100 µs/phase). The current amplitude used was empirically determined by delivering a short burst of stimulation with a current amplitude starting at 200 µA and increasing in 25 µA steps until we observed an obvious ElectraOFF response but not any behavioral changes. Electrical stimulation delivery was triggered using a MATLAB generated TTL pulse for 60 s, which started 15 s after triggering the first camera frame.

### Flicker Presentation

Visual flickers were presented via a 5 mm white LED (Amazon, MCIGICM) positioned in front of the animal. LED was pulsed at 4 Hz with a 50% duty cycle, for a total of 10 s, at an intensity of 20.5 µW in a dimly lit room. LED was triggered using a NI-DAQ (National Instruments, BNC-2110 and PCIe-6323), 10 s after the first camera frame.

### ElectraOFF fluorescence pre-processing to obtain Vm traces

All data analysis was performed offline using custom MATLAB (2024a, Mathworks Inc.). ElectraOFF fluorescence Videos were motion-corrected using a pairwise rigid motion correction algorithm as described previously^19^. Briefly, all trials for a single experimental recording were first concatenated. The shift of each frame was identified by calculating the maximum cross-correlation coefficient between each frame and the reference image. Regions of interest (ROIs) corresponding to individual neurons were subsequently manually identified using a max-minus-min projection image of the motion corrected data and the drawPolygon function (MATLAB). The ElectraOFF fluorescence trace was extracted by averaging all the pixels within the ROI to obtain the ROI trace. We then estimate the background trace for each ROI trace by averaging across all pixels of a manually selected area within the same imaging field of view, free of neuropil. The correlation coefficients of the background trace and the ROI trace were then obtained using the linear fit (y=mx+b), where m and b were found by minimizing the error between the background trace, x, and the ROI trace, y. The background trace is then scaled by plugging back in for x using the newly estimated coefficients to calculate a new y representing the portion of the ROI trace explainable by background fluctuations. ElectraOFF trace was then computed as the ROI trace minus the scaled background trace. ElectraOFF traces were further denoised by removing values greater than 10 standard deviations from the mean in the positive going direction. We then subtracted the median value of the trace, yielding the Vm trace for each neuron.

### Spike Identification, spike amplitude and spike-to-baseline ratio (SBR) calculation

To identify spikes, we first computed the baseline Vm fluctuations by averaging two traces yielded by (1) applying a moving average filter (window = 150 ms) of the Vm trace and, separately, (2) applying a second-order Savitsky-Golay filter (window = 150 ms) to the Vm trace. Baseline Vm fluctuations were then subtracted from the Vm trace to isolate the high frequency components (Vm_high_). Spikes were then detected using Vm_high_ as peaks greater than are 5 (CamKs) or 6 (ChIs and PVs) standard deviations away from the mean. Spike amplitude was calculated using the Vm trace as the difference between the spike peak and the minimum fluorescence value within the 3 frames (∼5 ms) preceding the spike peak. Identified spikes with an amplitude greater than 35 were considered artifacts and excluded from analysis. The SBR of each spike was computed as the spike amplitude divided by the approximated baseline noise. Baseline noise was defined for each trial as the standard deviation of the positive-half of Vm_high_ to exclude contributions from the negative-going spikes and slow baseline Vm fluctuations.

### Photostability Analysis

To assess the photostability of ElectraOFF *in vivo* we measured four parameters over time (1) mean ElectraOff raw fluorescence (F) for each 30 s trial, (2) normalized spike amplitude (dF/F) for each spike, (3) spike SBR for each spike, and (4) spike rate calculated for each 10 s bin.

Because the excitation LED power was occasionally adjusted to optimize recording length, raw F, dF/F and SBR changes were computed for each period of constant LED power. Within each period with constant LED power, data were linearly fit across time, and the percent-change per minute was calculated as the slope of the fit line divided by the starting y-value of the fit. Since spike rate was independent of LED power, changes in spike rate were estimated by linearly fitting over the full length of the recording.

### Spike triggered Vm Analysis

The spike-triggered Vm waveform for a neuron was calculated by first taking a 1-second window centered around each detected spike. Each window was scaled by its spike height, and then averaged across all spikes for the neuron. For visualization, final traces were flipped to follow the convention that spikes are positive-going.

When calculating the spike-triggered average Vm power spectra, spikes were first removed by replacing the 5 data points centered around each spike peak with linearly interpolated values to minimize broadband power changes at spike peaks. The power spectrum of the spike removed trace was then obtained for each 30 s trial using Matlab’s continuous wavelet transform function (cwt) (Morlet Wavelet, VoicesPerOctave = 12). The spectral trace was then windowed around each spike (1 s windows) and averaged across all spikes to obtain the spike triggered average Vm power spectra for a neuron.

### Visual flicker and electrical stimulation evoked responses

To assess the visual flicker and electrical stimulation evoked responses, Vm traces were first flipped, normalized to spike height, detrended, referenced relative to the median baseline Vm, and averaged across neurons. Briefly, we first obtained a scaled trace by normalizing every flipped trial trace to its average spike height. The scaled trace was then lowpass filtered (fifth order butterworth with 50 Hz cutoff) and fit with a linear trendline. This linear trendline was then subtracted from the scaled trace to account for photobleaching. A trial average trace was then calculated for each neuron by averaging these detrended traces. The median of the baseline period was then subtracted from the average trace, and the traces were then smoothed for visualization with a moving median filter (150 ms; 100 frames). The final reported mean trace and standard error were calculated using these smoothed trial-averaged traces.

To assess the effect of neuromodulation on firing rate, a binarized spike raster (each frame with a spike is 1) was generated for each trial, and the trial-average raster then calculated for each neuron. A moving mean filter (300 ms; 200 frames) was applied to calculate the firing rate trace. For each trial-average trace, the average baseline firing rate was calculated and subtracted to estimate the change in firing rate (Hz) relative to baseline, and the traces were then smoothed for visualization with a moving mean filter (150 ms; 100 frames) The final reported mean trace and standard error were calculated using these smoothed trial-averaged traces.

To classify neurons as entrained or not-entrained by 40 Hz electrical stimulation, each cell’s normalized trial traces were broken into 15 s chunks as seen in **Fig. 3c** and the power spectral density (PSD) of each of these chunks was calculated individually using Matlab’s fast-Fourier transform function (fft). The PSD around 40 Hz (39.5-40.51 Hz window) for each of the 15-second stimulation epochs was then Z-scored relative to the baseline PSD distribution in that frequency window. Neurons were classified as entrained if the normalized PSD at 40 Hz during the first 15 s of stimulation in any trial was greater than 3. Otherwise, neurons were classified as not-entrained.

## Acknowledgments

We thank members of Han Lab for their help throughout the study. We also thank Dr. Kiryl Piatkevich for providing the ElectraOFF plasmids and AAV9 particles. X. H. acknowledges funding from the NIH (1RF1NS129520, 1R01MH122971, and 1R01NS115797) and NSF (2002971-DIOS, and 1955981-CIF). C.R. acknowledges funding from NSF Graduate Research Fellowship 2021324226.

## Author contributions

Y.W., C.R. and X.H. designed the study. Y.W. collected experimental data.

C.R. analyzed the data. Y.Z. contributed some experimental data. Y.W., C.R. and X.H. prepared the manuscript, and all authors edited the manuscript. X.H. oversaw all aspects of the project and supervised the study.

